## Supplemental Appendix 1 for "Phosphorylation Toggles the SARS-CoV-2 Nucleocapsid Protein Between Two Membrane-Associated Condensate States"

**This PDF file includes:**

Table S1
Figures S1 to S18
Legends for Movies S1 to S6
SI References

**Other supporting materials for this manuscript include the following:**

Movies S1 to S6

**<u>Table S1</u>**

**Coding sequence for proteins, oligonucleotides, and recombinant DNA used in this study**

| Name | Sequence |
|---|---|
| SARS-COV-2 Nucleocapsid (N) protein | ATGCATCACCACCATCATCATGAGAATTTATATTTCCAGGGCAGCG<br>ATAACGGCCCGCAGAATCAGCGCAACGCGCCGCGCATTACCTTT<br>GGCGGCCCGAGCGATAGCACCGGCAGCAACCAAAACGGAGAGC<br>GCAGCGGCGCGCGCAGCAAACAACGCCGCCCCCAAGGCCTGCC<br>GAACAACACCGCGAGCTGGTTTACCGCGCTGACGCAGCATGGCA<br>AAGAAGATCTGAAATTTCCGCGCGGCCAAGGCGTGCCGATTAACA<br>CCAACAGCAGCCCGGATGATCAGATTGGCTATTATCGCCGCGCGA<br>CCCGCCGCATTCGCGGCGGCGATGGCAAAATGAAAGATCTGAGC<br>CCGCGCTGGTATTTTTATTATCTGGGCACCGGCCCGGAAGCGGG<br>CCTGCCGTATGGCGCGAACAAAGATGGCATTATTTGGGTGGCGAC<br>CGAAGGCGCGCTGAACACCCCGAAAGATCATATTGGCACCCGCA<br>ACCCGGCGAACAACGCGGCGATTGTGCTGCAGCTGCCGCAAGGC<br>ACCACCCTGCCGAAAGGCTTCTATGCCGAAGGTTCGCGCGGAGG<br>GAGCCAAGCGTCATCACGCAGCAGCAGCCGTTCGAGGAACAGCA<br>GCCGCAACAGCACTCCCGGGAGTTCCCGTGGCACATCGCCGGCG<br>CGTATGGCGGGCAATGGCGGGGACGCAGCGCTGGCGCTGCTGC<br>TGCTGGATCGCCTGAATCAGCTGGAAAGCAAAATGAGCGGCAAAG<br>GTCAGCAGCAGCAAGGCCAAACCGTAACGAAAAAAAGCGCGGCG<br>GAAGCGAGCAAAAAACCGCGTCAGAAACGCACCGCGACCAAAGC<br>GTATAACGTGACCCAAGCGTTTGGCCGCCGCGGCCCGGAACAGA<br>CCCAAGGCAACTTTGGCGATCAAGAACTGATTCGCCAAGGCACCG<br>ATTATAAACATTGGCCGCAGATTGCGCAGTTTGCGCCGAGCGCGA<br>GCGCGTTTTTTGGCATGAGCCGCATTGGCATGGAAGTGACCCCGA<br>GCGGCACCTGGCTGACCTATACCGGCGCGATTAAACTGGATGATA<br>AAGATCCGAACTTTAAAGATCAAGTGATTCTGCTGAACAAACATAT<br>TGATGCGTATAAAACCTTTCCGCCGACCGAACCGAAAAAAGATAA<br>AAAAAAAAAAGCGGATGAAACCCAAGCGCTGCCGCAGCGTCAGAA<br>AAAACAGCAGACCGTTACACTGCTGCCGGCGGCGGATCTGGATG<br>ATTTTAGCAAACAGCTGCAGCAGAGCATGAGCAGCGCGGATAGCA<br>CCCAAGCG |
| SL4 RNA FAM labeled | FAM – 5’<br>CUGUGUGGCUGUCACUCGGCUGCAUGCUUAGUGCACUCACGCA<br>G – 3’ |
| polyA RNA FAM labeled | FAM – 5’ AAAAAAAAAAAAAAAAAAAAAAAAAAAAAA – 3’ |
| 1-1000 RNA | GGGTTAAAGGTTTATACCTTCCCAGGTAACAAACCAACCAACTTTC<br>GATCTCTTGTAGATCTGTTCTCTAAACGAACTTTAAAATCTGTGTG<br>GCTGTCACTCGGCTGCATGCTTAGTGCACTCACGCAGTATAATTAA<br>TAACTAATTACTGTCGTTGACAGGACACGAGTAACTCGTCTATCTT<br>CTGCAGGCTGCTTACGGTTTCGTCCGTGTTGCAGCCGATCATCAG<br>CACATCTAGGTTTCGTCCGGGTGTGACCGAAAGGTAAGATGGAGA<br>GCCTTGTCCCTGGTTTCAACGAGAAAACACACGTCCAACTCAGTTT |

| | |
|---|---|
| | GCCTGTTTTACAGGTTCGCGACGTGCTCGTACGTGGCTTTGGAGA<br>CTCCGTGGAGGAGGTCTTATCAGAGGCACGTCAACATCTTAAAGA<br>TGGCACTTGTGGCTTAGTAGAAGTTGAAAAAGGCGTTTTGCCTCAA<br>CTTGAACAGCCCTATGTGTTCATCAAACGTTCGGATGCTCGAACTG<br>CACCTCATGGTCATGTTATGGTTGAGCTGGTAGCAGAACTCGAAG<br>GCATTCAGTACGGTCGTAGTGGTGAGACACTTGGTGTCCTTGTCC<br>CTCATGTGGGCGAAATACCAGTGGCTTACCGCAAGGTTCTTCTTC<br>GTAAGAACGGTAATAAAGGAGCTGGTGGCCATAGTTACGGCGCC<br>GATCTAAAGTCATTTGACTTAGGCGACGAGCTTGGCACTGATCCTT<br>ATGAAGATTTTCAAGAAAACTGGAACACTAAACATAGCAGTGGTGT<br>TACCCGTGAACTCATGCGTGAGCTTAACGGAGGGGCATACACTCG<br>CTATGTCGATAACAACTTCTGTGGCCCTGATGGCTACCCTCTTGAG<br>TGCATTAAAGACCTTCTAGCACGTGCTGGTAAAGCTTCATGCACTT<br>TGTCCGAACAACTGGACTTTATTGACACTAAGAGGGGTGTATACT<br>GCTGCCGTGAACATGAGCATGAAATTGCTTGGTACACGGAACGTT<br>CTGGGCCCTCGA |
| N RNA | GGGTTAAAGGTTTATACCTTCCCAGGTAACAAACCAACCAACTTTC<br>GATCTCTTGTAGATCTGTTCTCTAAACGAACAAACTAAAATGTCTG<br>ATAATGGACCCCAAAATCAGCGAAATGCACCCCGCATTACGTTTG<br>GTGGACCCTCAGATTCAACTGGCAGTAACCAGAATGGAGAACGCA<br>GTGGGGCGCGATCAAAACAACGTCGGCCCCAAGGTTTACCCAATA<br>ATACTGCGTCTTGGTTCACCGCTCTCACTCAACATGGCAAGGAAG<br>ACCTTAAATTCCCTCGAGGACAAGGCGTTCCAATTAACACCAATAG<br>CAGTCCAGATGACCAAATTGGCTACTACCGAAGAGCTACCAGACG<br>AATTCGTGGTGGTGACGGTAAAATGAAAGATCTCAGTCCAAGATG<br>GTATTTCTACTACCTAGGAACTGGGCCAGAAGCTGGACTTCCCTAT<br>GGTGCTAACAAAGACGGCATCATATGGGTTGCAACTGAGGGAGC<br>CTTGAATACACCAAAAGATCACATTGGCACCCGCAATCCTGCTAAC<br>AATGCTGCAATCGTGCTACAACTTCCTCAAGGAACAACATTGCCAA<br>AAGGCTTCTACGCAGAAGGGAGCAGAGGCGGCAGTCAAGCCTCT<br>TCTCGTTCCTCATCACGTAGTCGCAACAGTTCAAGAAATTCAACTC<br>CAGGCAGCAGTAGGGGAACTTCTCCTGCTAGAATGGCTGGCAAT<br>GGCGGTGATGCTGCTCTTGCTTTGCTGCTGCTTGACAGATTGAAC<br>CAGCTTGAGAGCAAAATGTCTGGTAAAGGCCAACAACAACAAGGC<br>CAAACTGTCACTAAGAAATCTGCTGCTGAGGCTTCTAAGAAGCCT<br>CGGCAAAAACGTACTGCCACTAAAGCATACAATGTAACACAAGCTT<br>TCGGCAGACGTGGTCCAGAACAAACCCAAGGAAATTTTGGGGACC<br>AGGAACTAATCAGACAAGGAACTGATTACAAACATTGGCCGCAAA<br>TTGCACAATTTGCCCCCAGCGCTTCAGCGTTCTTCGGAATGTCGC<br>GCATTGGCATGGAAGTCACACCTTCGGGAACGTGGTTGACCTACA<br>CAGGTGCCATCAAATTGGATGACAAAGATCCAAATTTCAAAGATCA<br>AGTCATTTTGCTGAATAAGCATATTGACGCATACAAAACATTCCCA<br>CCAACAGAGCCTAAAAAGGACAAAAAGAAGAAGGCTGATGAAACT<br>CAAGCCTTACCGCAGAGACAGAAGAAACAGCAAACTGTGACTCTT<br>CTTCCTGCTGCAGATTTGGATGATTTCTCCAAACAATTGCAACAAT<br>CCATGAGCAGTGCTGACTCAACTCAGG |
| Primer 1-1000 Forward | CCATCCGGCGTAATACGACTCACTATAGGG |

| | |
|---|---|
| Primer 1-1000 Reverse | CTAGAAAGATAGAACGTTCCGTGTACCAAG |
| Primer N Forward | GTGTGATGGATATCTGCAGAATTCGC |
| Primer N Reverse | CATGAGTTTAGGCCTGAGTTGAGTCAG |
| 6xHis-GFP-M(104-222) | ATGGGTTCTTCTCACCATCACCATCACCATGGTTCTTCTGTGAGCA AGGGCGAGGAGCTGTTCACCGGGGTGGTGCCCATCCTGGTCGAG CTGGACGGCGACGTAAACGGCCACAAGTTCAGCGTGCGCGGCGA GGGCGAGGGCGATGCCACCAACGGCAAGCTGACCCTGAAGTTCA TCTGCACCACCGGCAAGCTGCCCGTGCCCTGGCCCACCCTCGTG ACCACCCTGACCTACGGCGTGCAGTGCTTCAGCCGCTACCCCGA CCACATGAAGCAGCACGACTTCTTCAAGTCCGCCATGCCCGAAGG CTACGTCCAGGAGCGCACCATCTCCTTCAAGGACGACGGCACCTA CAAGACCCGCGCCGAGGTGAAGTTCGAGGGCGACACCCTGGTGA ACCGCATCGAGCTGAAGGGCATCGACTTCAAGGAGGACGGCAAC ATCCTGGGGCACAAGCTGGAGTACAACTTCAACAGCCACAACGTC TATATCACGGCCGACAAGCAGAAGAACGGCATCAAGGCGAACTTC AAGATCCGCCACAACGTCGAGGACGGCAGCGTGCAGCTCGCCGA CCACTACCAGCAGAACACCCCCATCGGCGACGGCCCCGTGCTGC TGCCCGACAACCACTACCTGAGCACCCAGTCCAAGCTGAGCAAAG ACCCCAACGAGAAGCGCGATCACATGGTCCTGCTGGAGTTCGTG ACCGCCGCCGGGATCACTCTCGGCATGGACGAGCTGTACAAGGG GATCGAGGAAAACCTGTACTTCCAATCCAATGCAGCTCGCACACG CAGTATGTGGTCCTTTAACCCGGAGACCAATATTCTTCTGAACGTC CCCTTGCATGGTACTATCCTTACCCGCCCCCTTCTGGAGAGTGAA CTGGTGATCGGTGCCGTCATCTTACGTGGGCATTTACGCATCGCG GGGCACCACTTAGGGCGCTGTGACATTAAAGACTTACCCAAGGAA ATTACTGTAGCTACTTCGCGTACTCTTTCCTATTATAAGTTAGGCG CATCACAGCGCGTGGCGGGCGATTCTGGCTTTGCAGCATATTCAC GCTACCGCATTGGGAATTATAAATTAAATACAGATCACTCAAGTTC CTCCGATAACATCGCCCTGTTGGTACAG |
| 6xHis-GFP-Nsp3Ubl1 | ATGGGTTCTTCTCACCATCACCATCACCATGGTTCTTCTGTGAGCA AGGGCGAGGAGCTGTTCACCGGGGTGGTGCCCATCCTGGTCGAG CTGGACGGCGACGTAAACGGCCACAAGTTCAGCGTGCGCGGCGA GGGCGAGGGCGATGCCACCAACGGCAAGCTGACCCTGAAGTTCA TCTGCACCACCGGCAAGCTGCCCGTGCCCTGGCCCACCCTCGTG ACCACCCTGACCTACGGCGTGCAGTGCTTCAGCCGCTACCCCGA CCACATGAAGCAGCACGACTTCTTCAAGTCCGCCATGCCCGAAGG CTACGTCCAGGAGCGCACCATCTCCTTCAAGGACGACGGCACCTA CAAGACCCGCGCCGAGGTGAAGTTCGAGGGCGACACCCTGGTGA ACCGCATCGAGCTGAAGGGCATCGACTTCAAGGAGGACGGCAAC ATCCTGGGGCACAAGCTGGAGTACAACTTCAACAGCCACAACGTC TATATCACGGCCGACAAGCAGAAGAACGGCATCAAGGCGAACTTC AAGATCCGCCACAACGTCGAGGACGGCAGCGTGCAGCTCGCCGA CCACTACCAGCAGAACACCCCCATCGGCGACGGCCCCGTGCTGC TGCCCGACAACCACTACCTGAGCACCCAGTCCAAGCTGAGCAAAG ACCCCAACGAGAAGCGCGATCACATGGTCCTGCTGGAGTTCGTG ACCGCCGCCGGGATCACTCTCGGCATGGACGAGCTGTACAAGGG |

| | |
|---|---|
| | GATCGAGGAAAACCTGTACTTCCAATCCAATGCATCTTCTAATGGC<br>GCACCGACAAAAGTTACATTTGGAGACGATACCGTGATCGAAGTT<br>CAGGGCTACAAAAGCGTGAACATCACCTTCGAGCTGGATGAACGT<br>ATCGATAAAGTGCTGAACGAGAAATGCAGCGCATATACCGTGGAA<br>CTGGGTACCGAAGTGAACGAATTTGCCTGTGTTGTTGCAGATGCA<br>GTGATCAAAACCTTACAGCCGGTTAGCGAACTGCTGACACCTTTA<br>GGCATTGATCTGGATGAATGGAGCATGGCAACCTATTATCTGTTC<br>GACGAAAGCGGCGAGTTCAAACTGGCATCACACATGTATTGCAGC<br>TTCTATCCGCCTGATGAA |

**Supplementary Figures**

A. Unmodified N, theoretical mass: 47300 Da

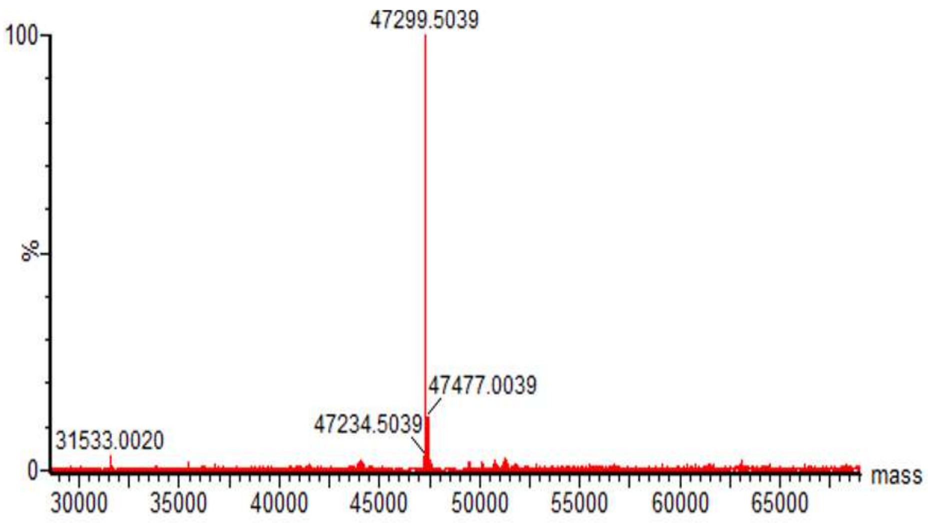


Phosphorylated N, theoretical mass with 9 phosphate groups: 48020 Da

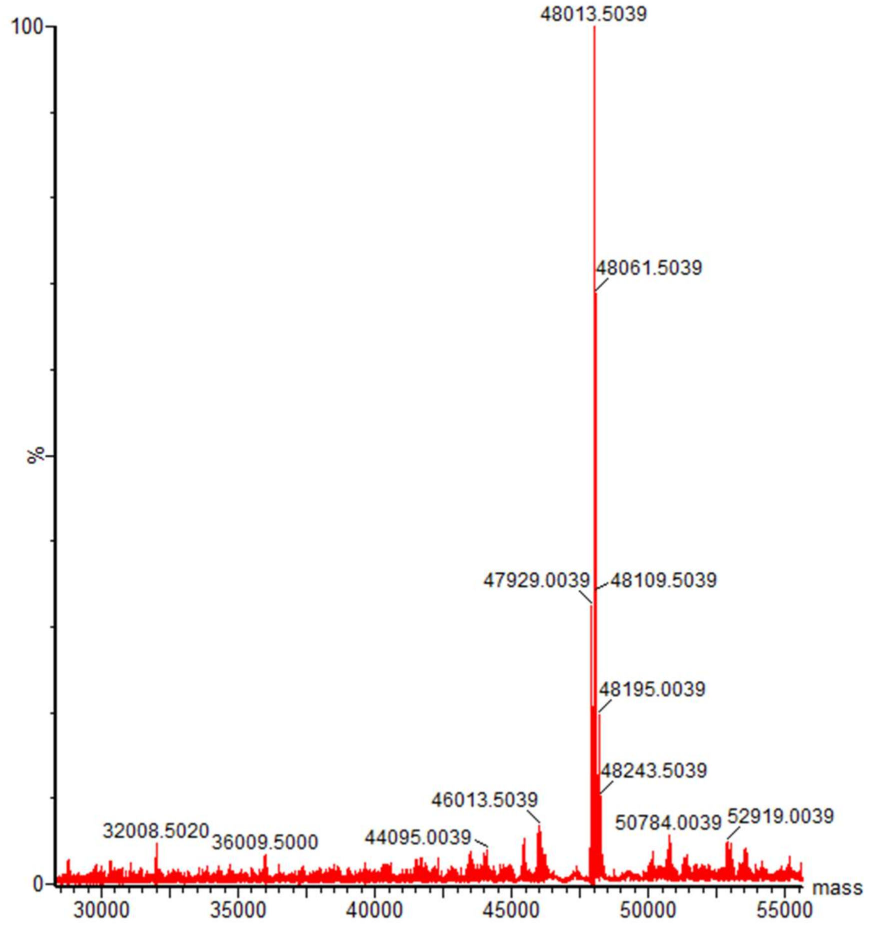

**B.**

Peptide: 172 AEGSRGGSQASSRSSSRSRNSSRNSTPGSSRGTSPARMAGNGGDAAL 219

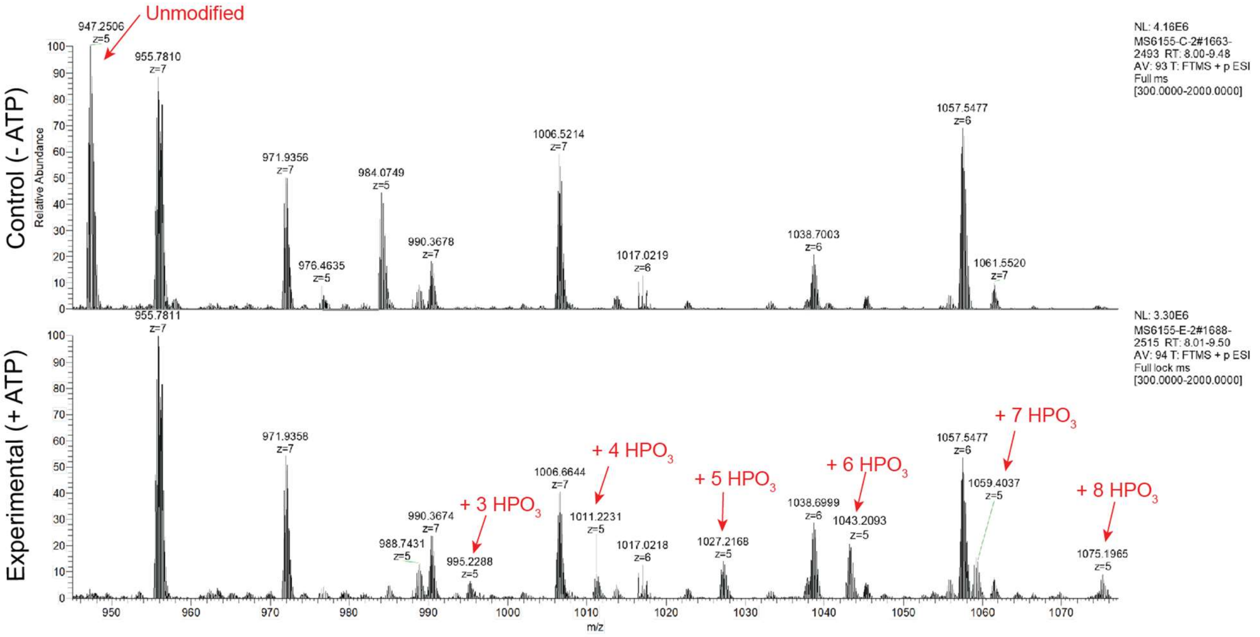


**C.** Peptide: 170 GFYAEG**S**R 177

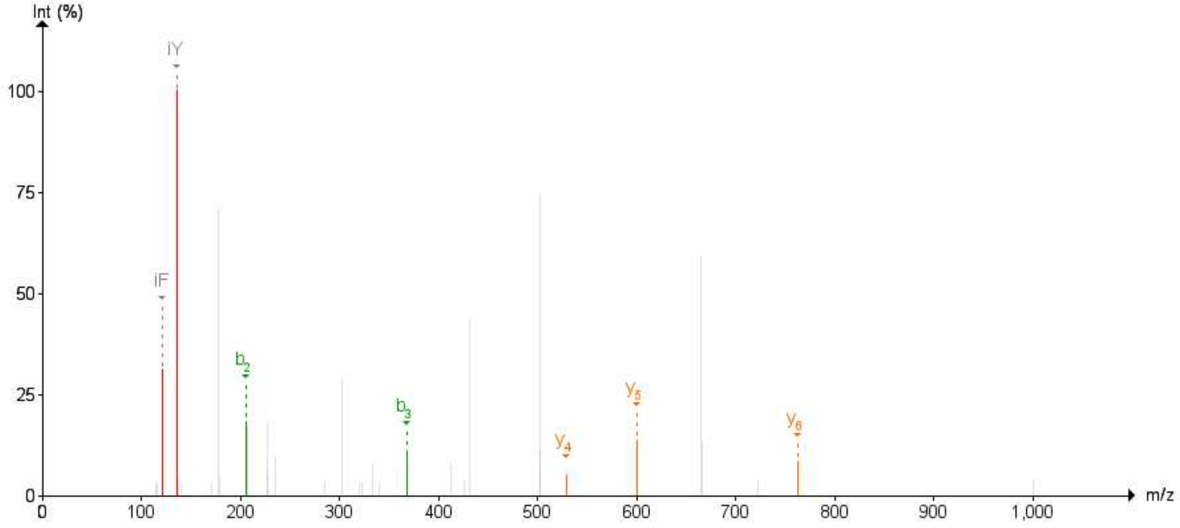

Peptide: 170 GFYAEG**S**RGG**S**QAS**S**R 185

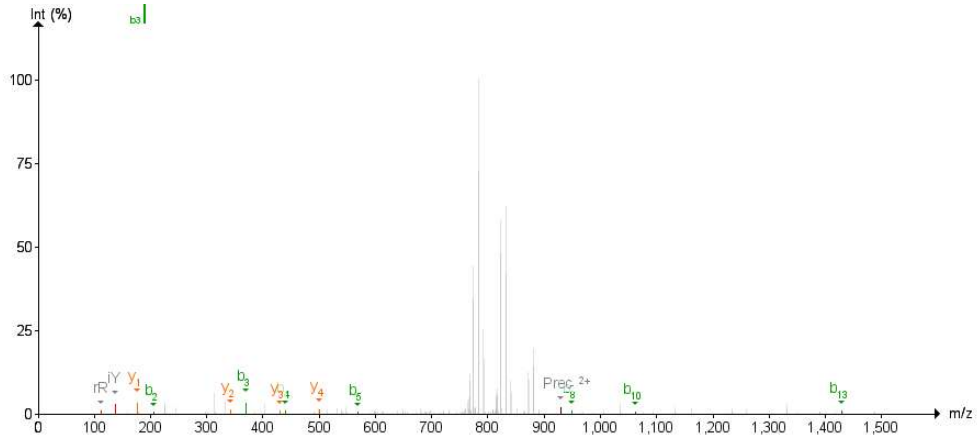


Peptide: 170 GFYAEG**S**RGG**S**QAS**S**R 185

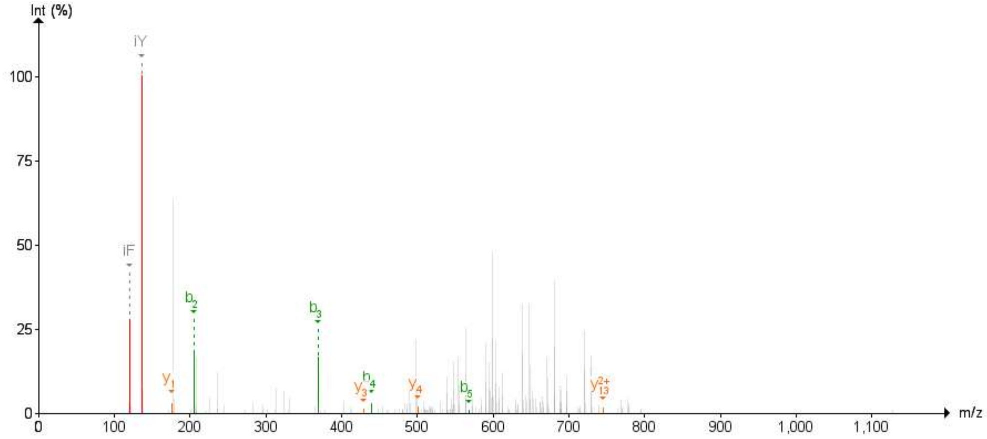


Peptide: 196 NS**T**PGS**S**RGT**S**PAR 209

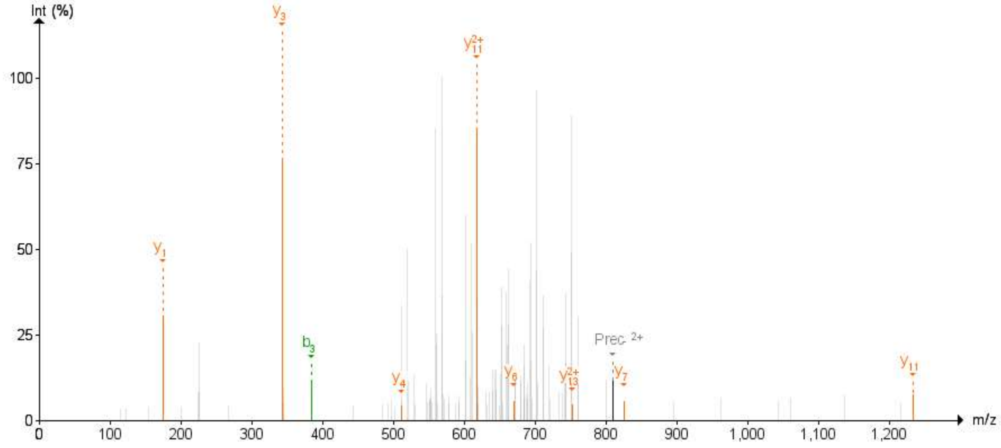

Peptide: 196 NS**T**PGS**S**RGT**S**PAR 209

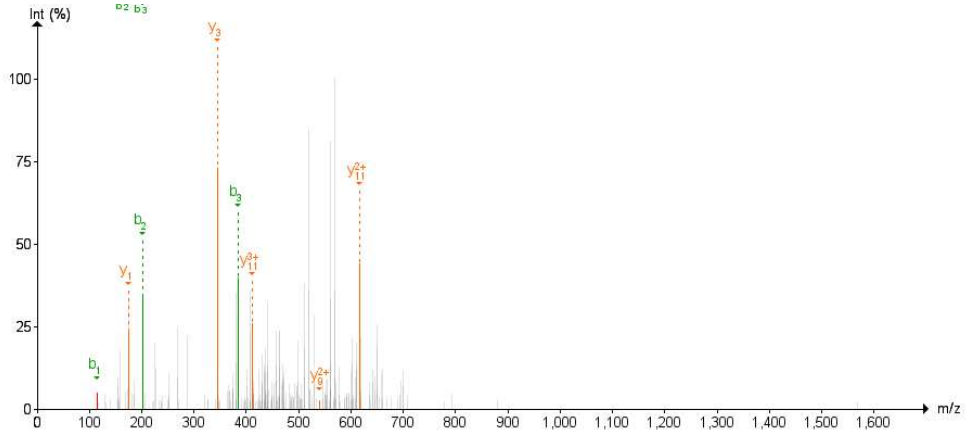


**D.** Phos-tag gel electrophoresis

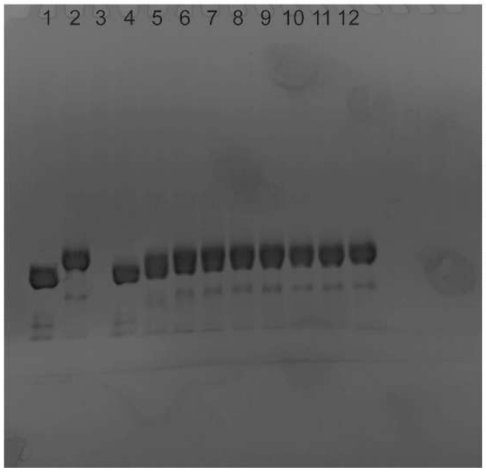


1. N
2. N + SRPK + GSK3β + Rxn buffer + 120 min
3. Empty
4. Rxn mixture at 0 min
5. Rxn mixture at 15 min
6. Rxn mixture at 30 min
7. Rxn mixture at 45 min
8. Rxn mixture at 60 min
9. Rxn mixture at 75 min
10. Rxn mixture at 90 min
11. Rxn mixture at 105 min
12. Rxn mixture at 120 min

**Figure S1. Confirmation of N protein phosphorylation**. A) Mass spectra for N protein prior to phosphorylation and following phosphorylation, showing a mass increase of the main peak from 47.3 kDa to 48.0 kDa. This mass increase represents the addition of 9 (+ 720 Da) phosphate groups. B) Following digestion of N / pN protein with pepsin protease, we conducted LC-MS on the peptide fragments. (Top) Mass spectra of peptides from a control sample (N + SRPK1 + GSK3β without ATP). A peptide containing the SR-rich region (residues 172-219) was identified and labelled as "unmodified" (with sequence noted above). (Bottom) Mass spectra of peptides from a sample with phosphorylated N (N + SRPK1 + GSK3β + ATP). The peak representing the unmodified peptide can no longer be identified. New peaks with the expected m/z of peptides with 3-8 phosphate groups added to the peptide can be identified and are labelled. The presence of peptides with a range of phosphorylation states indicates variability in the number of sites that are successfully phosphorylated per protein. C) Representative mass spectra identifying phosphorylation sites in peptides from pN protein. Following digestion of pN with trypsin protease, we conducted LC-MS on the peptide fragments. 5 peptides were identified from which phosphorylation sites could be precisely identified: S176, S180, S184, S184, T198, S202, S206, which are highlighted in bold above the respective spectra. Additional phosphorylation sites could be detected in other peptides, but the precise site of phosphorylation could not be identified due to the protease digestion patterns. All sites had been previously identified by Yaron et al., 2022. D) SuperSep Phos-tag SDS-PAGE of N protein sample before, after, and during phosphorylation reaction at the timepoints indicated.

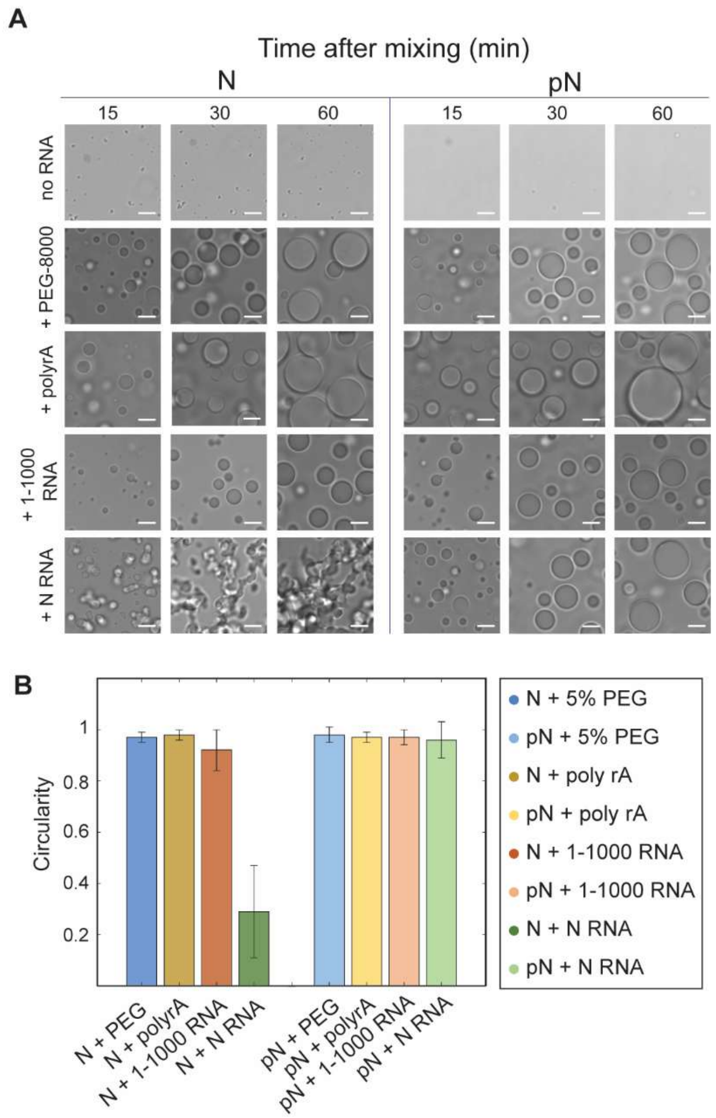


**Figure S2. Characterization of morphology of N condensate samples**. A) Representative images from the formation of condensates over time with N or pN protein – without RNA, with a crowding agent (5% PEG8000), with unstructured polyA RNA or with viral RNA fragments. Droplet morphology depends on protein and RNA combination. Scale bar = 5 μm. B) Quantification of droplet morphology from A. Circularity of droplets at 60 minutes was measured as:

$$\boldsymbol{C = (4\,\pi * area/perimeter^2) * (1 - 0.5/r)^2}$$

where $\boldsymbol{r} = \boldsymbol{perimeter}/2\boldsymbol{\pi}$. All protein and RNA combinations have a circularity ~ 1 except for condensates formed from N protein and N RNA. (n = 5 independent trials).

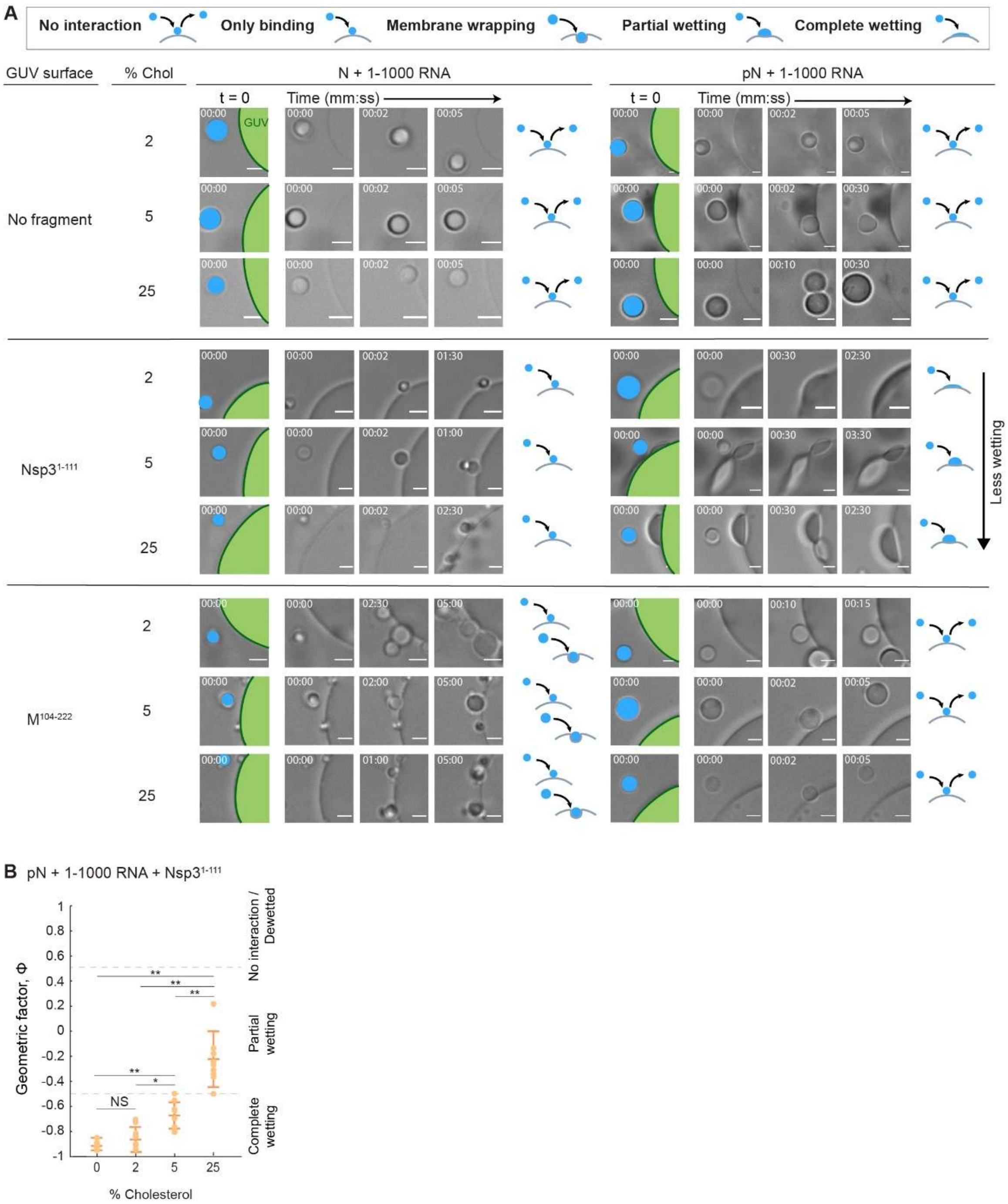


**Figure S3. Effect of cholesterol on N condensate interaction with membranes**. A) Representative widefield images showing the interaction between condensates and membranes over time, with GUVs incorporating 2, 5, and 25% cholesterol, resulting in final lipid compositions of:

+ 2% cholesterol: 58% DOPC, 25% DOPE, 5% Ni-NTA, 10% DOPS, 2% cholesterol

+ 5% cholesterol: 55% DOPC, 25% DOPE, 5% Ni-NTA, 10% DOPS, 5% cholesterol

+ 25% cholesterol: 35% DOPC, 25% DOPE, 5% Ni-NTA, 10% DOPS, 25% cholesterol. GUVs are labeled in green, and condensates in blue. Interaction type is qualitatively classified. Scale bars = 5 μm. B) Quantification of the geometric factor from the interaction of pN and 1-1000 RNA condensates, with GUVs coated with $NSP3^{1-111}$ and varying cholesterol compositions (data with 0% cholesterol from Figure 2). p values were determined using one-way ANOVA followed by post hoc Tukey's test. (NS, not significant; * $p < 0.05$; ** $p < 0.01$).

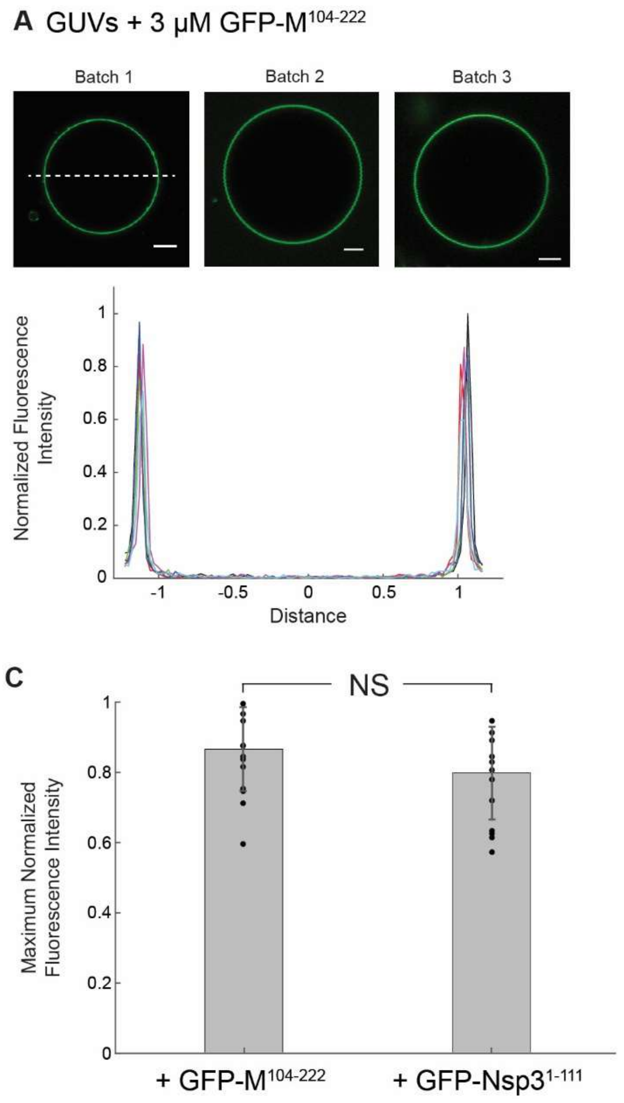


**Figure S4. Degree of tethering of membrane protein fragments to the GUV surface.** A) (top) Representative confocal images from three GUV batches with 3 μM GFP-$M^{104-222}$ added to the sample. Fluorescence intensities were quantified as line profiles across individual GUVs as indicated by the example dashed white line. Scale bar = 5 μm. (bottom) Fluorescence intensity profiles across one GUV from each batch made (n = 6), normalized by GUV size. Intensities were also divided by $2^{16}$, the dynamic range of the images, such that intensities scale from 0 to 1. B) (top) Representative confocal images from three GUV batches with 3 μM GFP-Nsp3$^{1-111}$ added to the sample. Scale bar = 5 μm. (bottom) Fluorescence intensity profiles across one GUV from each batch made (n = 6), normalized by GUV size. Intensities were also divided by $2^{16}$, the dynamic range of the images, such that intensities scale from 0 to 1. C) Maximum fluorescence intensity measured in the GFP-M$^{104-222}$ and GFP-Nsp3$^{1-111}$ samples. The difference in intensity is not statistically significant ($p = 0.2$). p value was determined using a two-sided student's t-test.

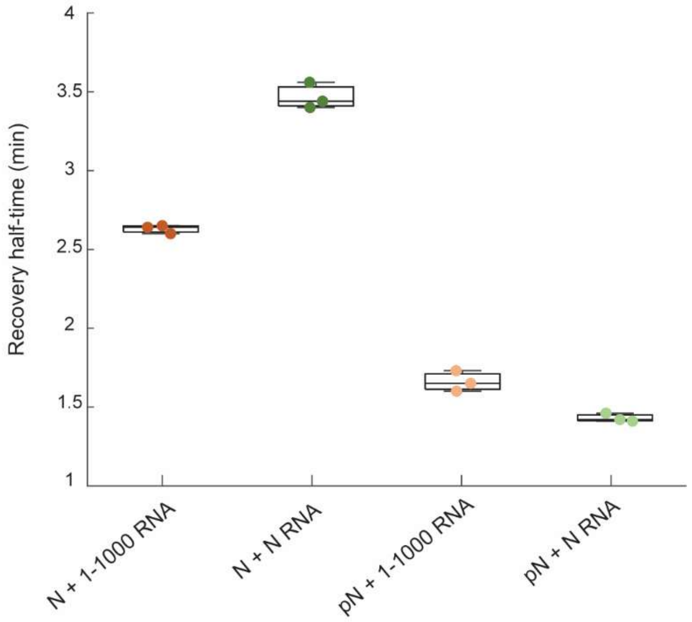


**Figure S5. Recovery half times quantified from fluorescence recovery after photobleaching (FRAP) experiments.** FRAP recovery curves following protein bleaching were fit to a simple exponential model. Phosphorylation reduces the recovery half time of N protein in condensates, indicating an increase in the diffusion of N. Data represents 3 independent trials.

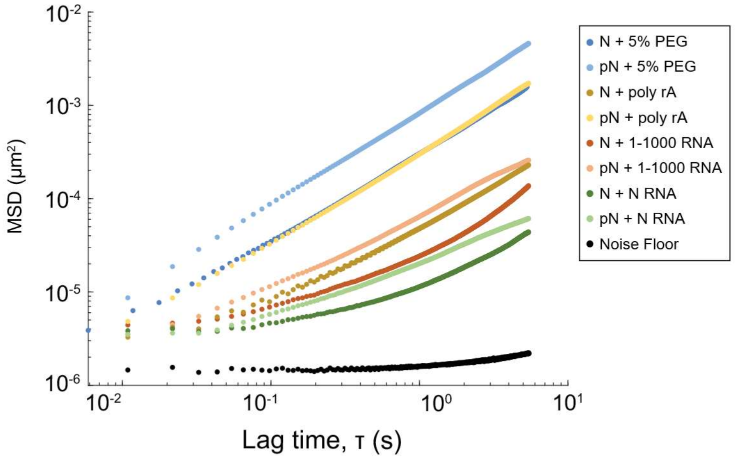


**Figure S6. Noise floor for the MSD data shown in Figure 4D.** Ensemble MSD versus lag time for the protein and RNA combinations tested in this study, including the noise floor in black. The noise floor was calculated as the average of 12 videos from 3 independent trials. The experimental setup includes a microfluidic temperature controller (Cherry Biotech) that causes vibrations resulting in noise.

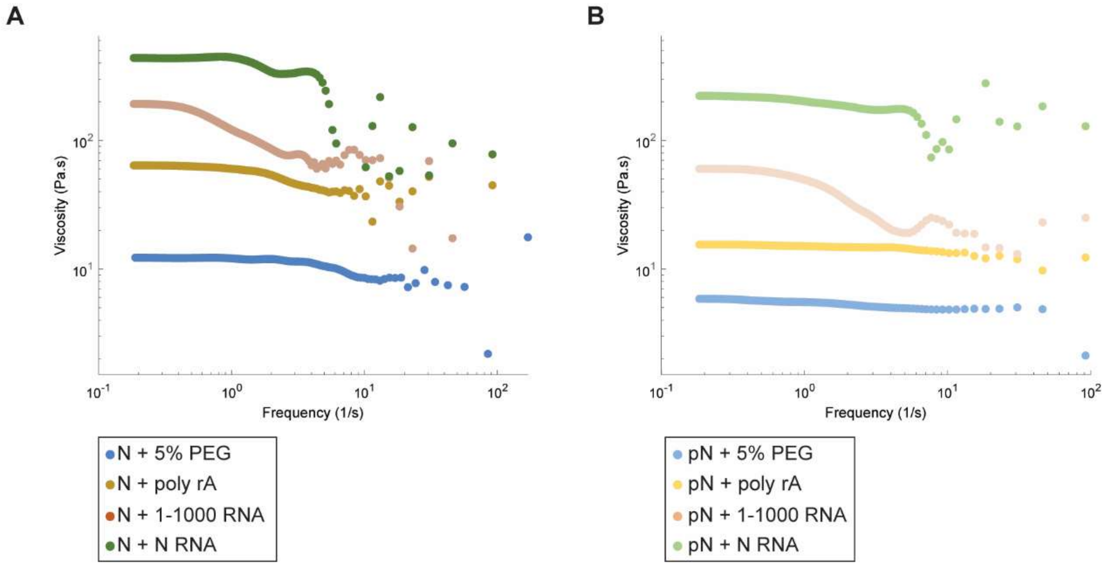


**Figure S7. The viscosity of N condensates as a function of frequency as obtained from the microrheology experiments.** These results were used to calculate the zero-shear viscosities plotted in Figure 4 and S10.

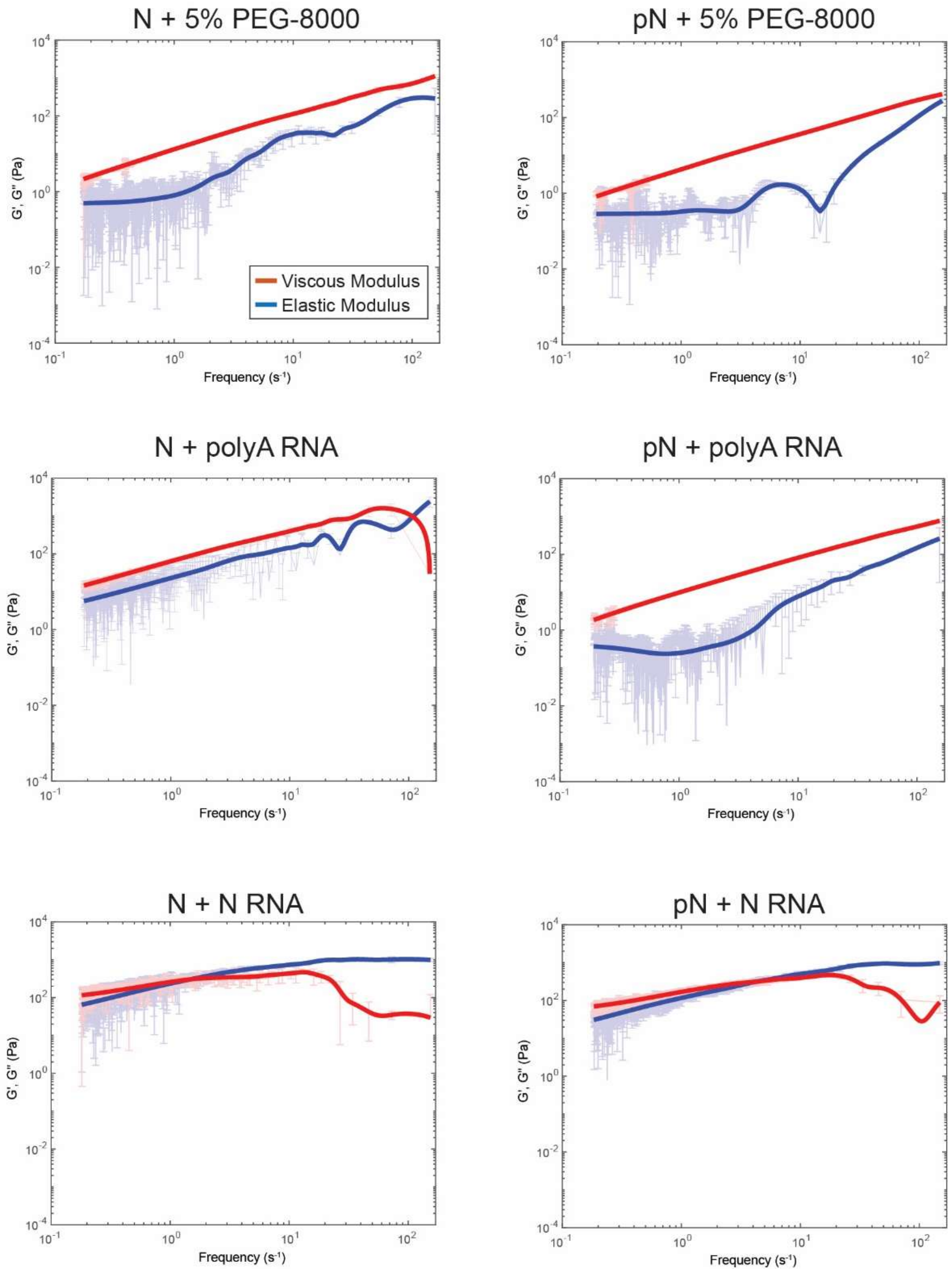


**Figure S8. Frequency-dependent viscous and elastic moduli for N vs. pN condensates not shown in Figure 4.** Plot with the average frequency-dependent viscous modulus (G'', red) and elastic modulus (G', blue) of N/pN + 5% PEG or N/pN +

1 mg/mL polyA RNA or N/pN + 300 nM N RNA condensates. Data is calculated from the MSDs in Figure S6 (from n ≥ 10 videos over 3 independent samples).

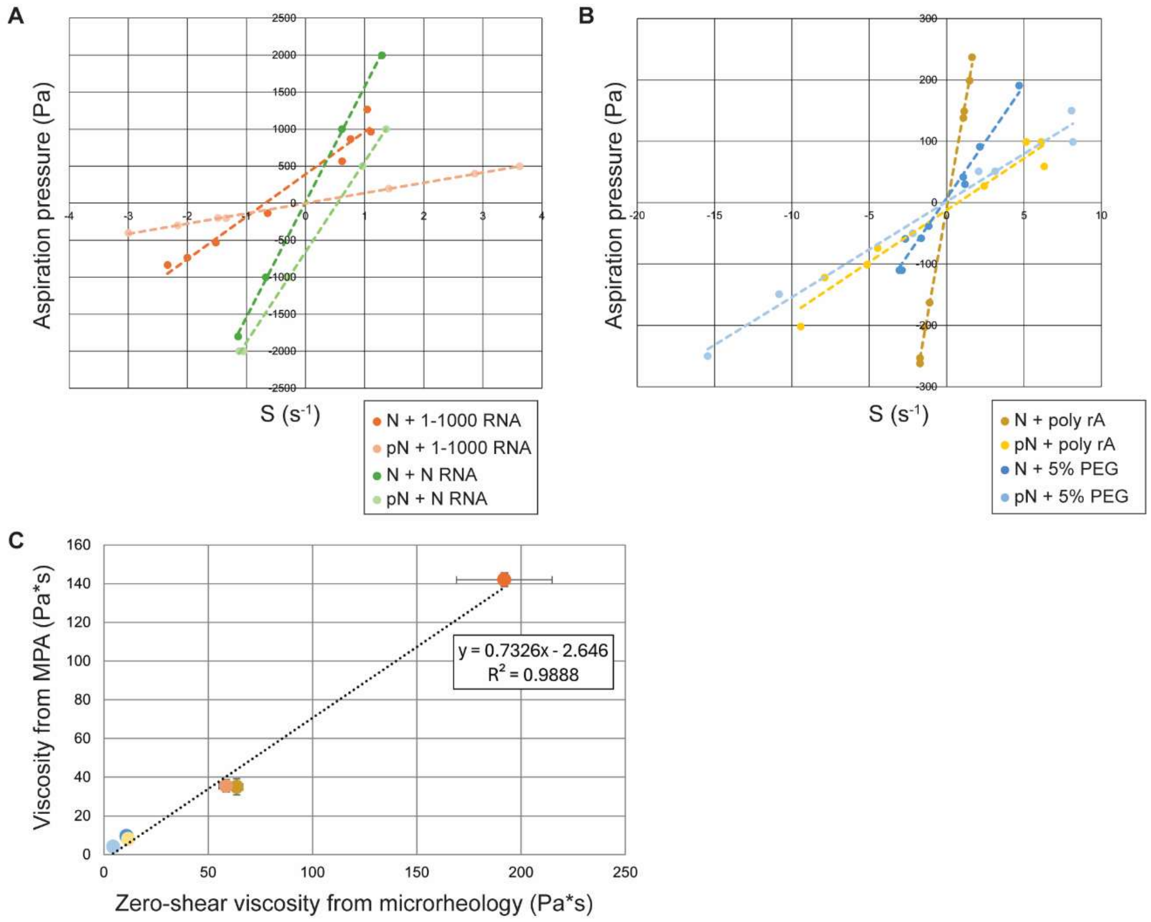


**Figure S9. Micropipette aspiration (MPA) of N/pN protein condensates.** A) The relationship between aspiration pressure and shear rate S, defined as $S = d(Lp/Rp)^2/dt$, where Lp is the aspiration length, Rp is the radius of the pipette, and t is time, for N or pN condensates with viral RNA. The viscosity is calculated as the slope of the best fit line divided by 4. Data is plotted in Figure 4I. B) Aspiration pressure vs. shear rate S for N or pN condensates with 5% PEG-8000 and 1 mg/mL polyA RNA. C) Comparison of viscosities measured for each N and pN condensate via MPA and microrheology. The trend in viscosities for the condensate compositions tested is consistent between the two methods, but MPA measurements were consistently lower than microrheology measurements. Plotted against each other, we obtain a slope of 0.72 and an $R^2$ fit of 0.99.

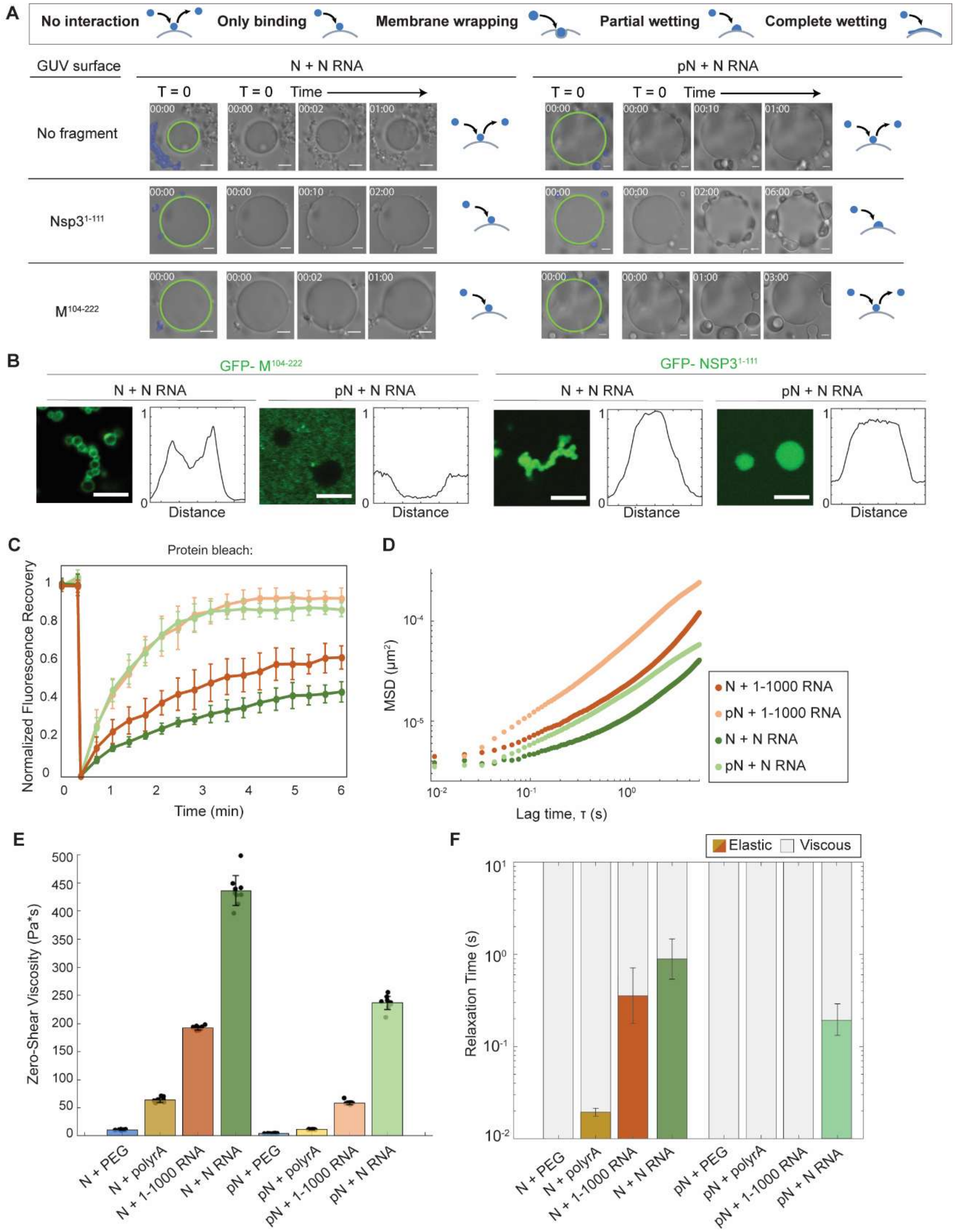


**Figure S10. Repetition of material property experiments with N RNA.** A) These experiments assess how condensates formed from N or pN protein plus N RNA interact with membranes. Using optical tweezers, condensates are trapped and brought to the surface of GUVs. GUVs have either no protein fragment or GFP-Nsp3$^{1-111}$ or GFP-M$^{104-222}$ displayed at their surface. Representative images show N or pN condensates with N RNA do not interact with naked membranes, but do bind to and may wet the surface of GUVs with Nsp3 depending on the N protein phosphorylation status. Condensates with N but not pN interact with GUVs with the M protein fragment. B) Interaction between N and membrane protein fragments was confirmed using a partitioning experiment. C) Fluorescence recovery after photobleaching with N/pN and N RNA compared to N/pN and 1-1000 RNA. Unmodified N has lower mobility when condensed with N RNA (recovery half-life = 3.5 ± 0.1 min), compared to with 1-1000 RNA (recovery half-life = 2.6 ± 0.1 min). pN recovery curves are similar across RNA samples (pN + 1-1000 RNA recovery half-life = 1.6 ± 0.1 min vs. 1.4 ± 0.1 for N RNA. D) Ensemble MSD versus lag time for N or pN and N RNA vs. 1-1000 RNA. E) The zero-shear viscosity of the protein and RNA condensates studied, calculated from the particle-tracking results after noise correction. Data from n ≥ 10 videos from 3 independent trials. F) Quantification of the timescales at which the elastic modulus dominates (color) versus the viscous modulus dominates (grey) in protein and RNA condensates. Error bars represent one standard deviation (±1 s.d.).

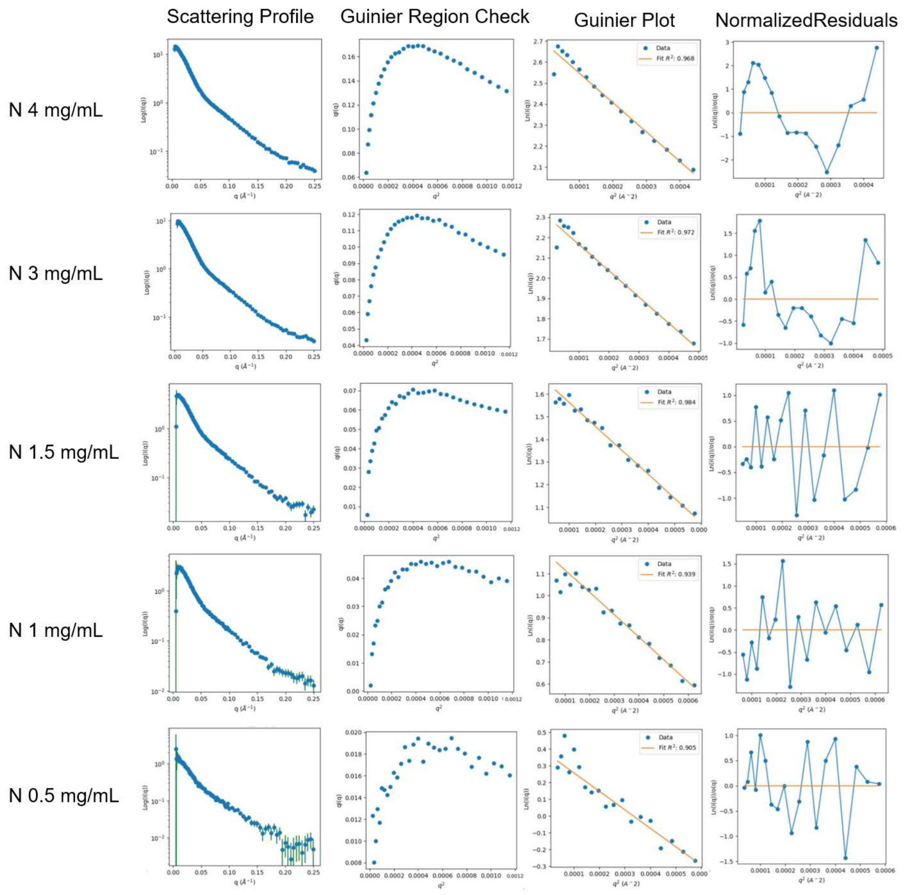


**Figure S11. Extended data for Small Angle X-Ray Scattering experiments for unmodified N protein at different concentrations.** First column, raw data in the form of scattering profiles. Second column, the Guinier peak analysis plot indicates the validity of doing the Guinier analysis in the third column[1]. Fourth column, analysis that the normalized residuals are randomly distributed about zero. Note, 4 mg/mL data was excluded from the analysis due to aggregation.

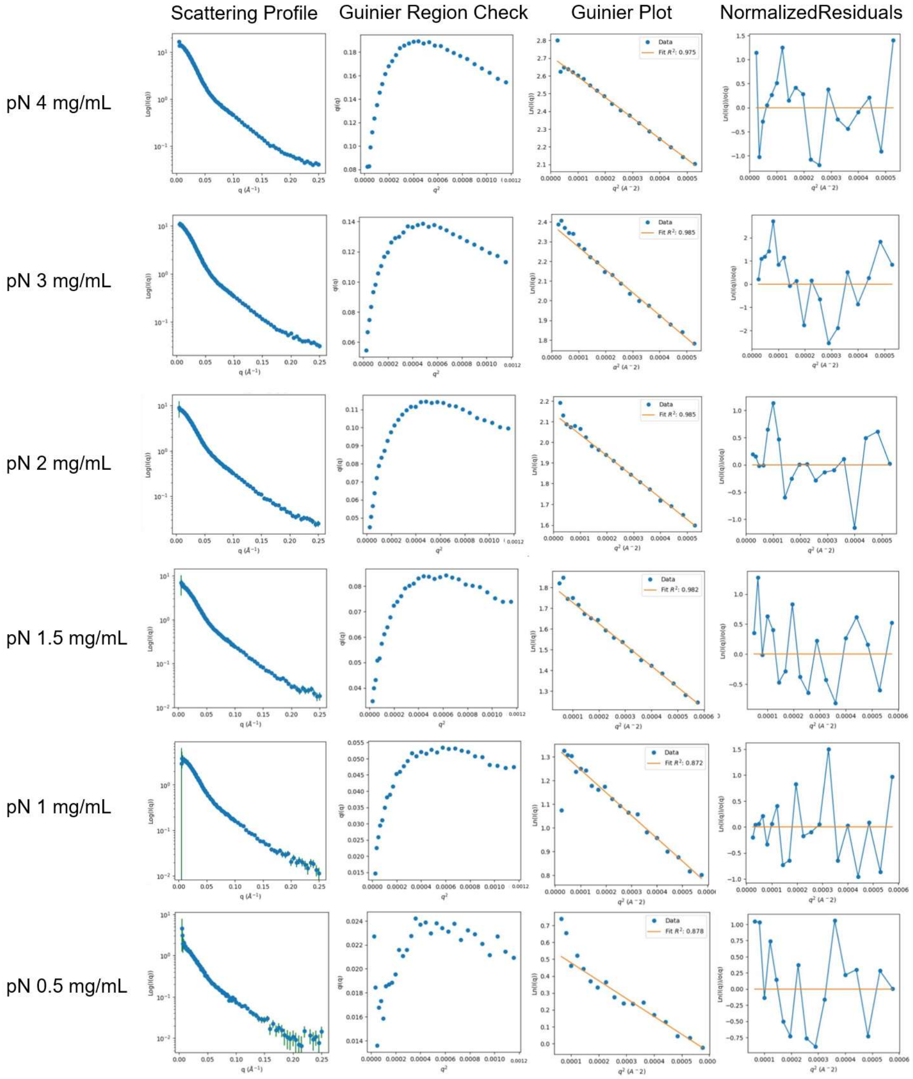


**Figure S12. Extended data for Small Angle X-Ray Scattering experiments for phosphorylated N protein at different concentrations.** First column, raw data in the form of scattering profiles. Second column, the Guinier peak analysis plot indicates the validity of doing the Guinier analysis in the third column[1]. Fourth column, analysis that the normalized residuals are randomly distributed about zero. Note, 4 mg/mL data was excluded from the analysis due to aggregation.

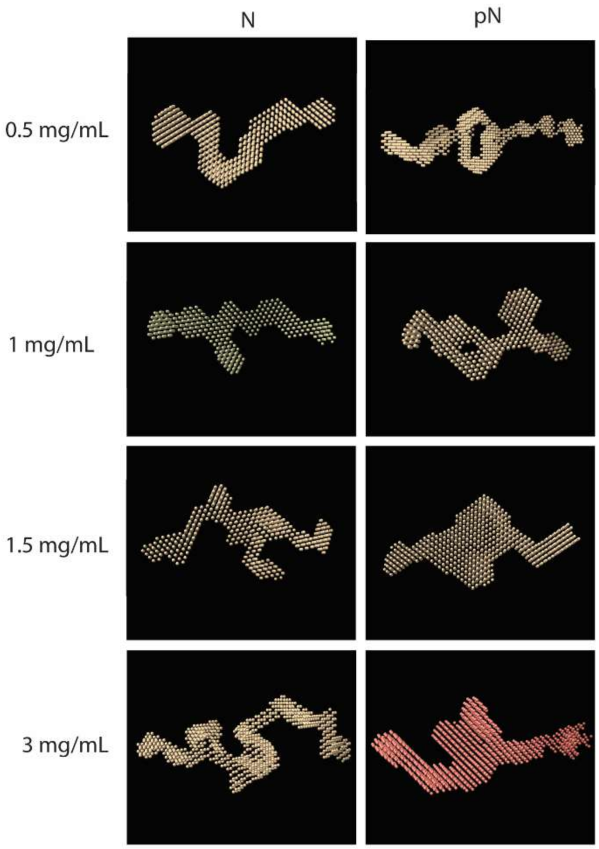


**Figure S13. Bead model representations for N and pN from SAXS data at varying protein concentrations.**

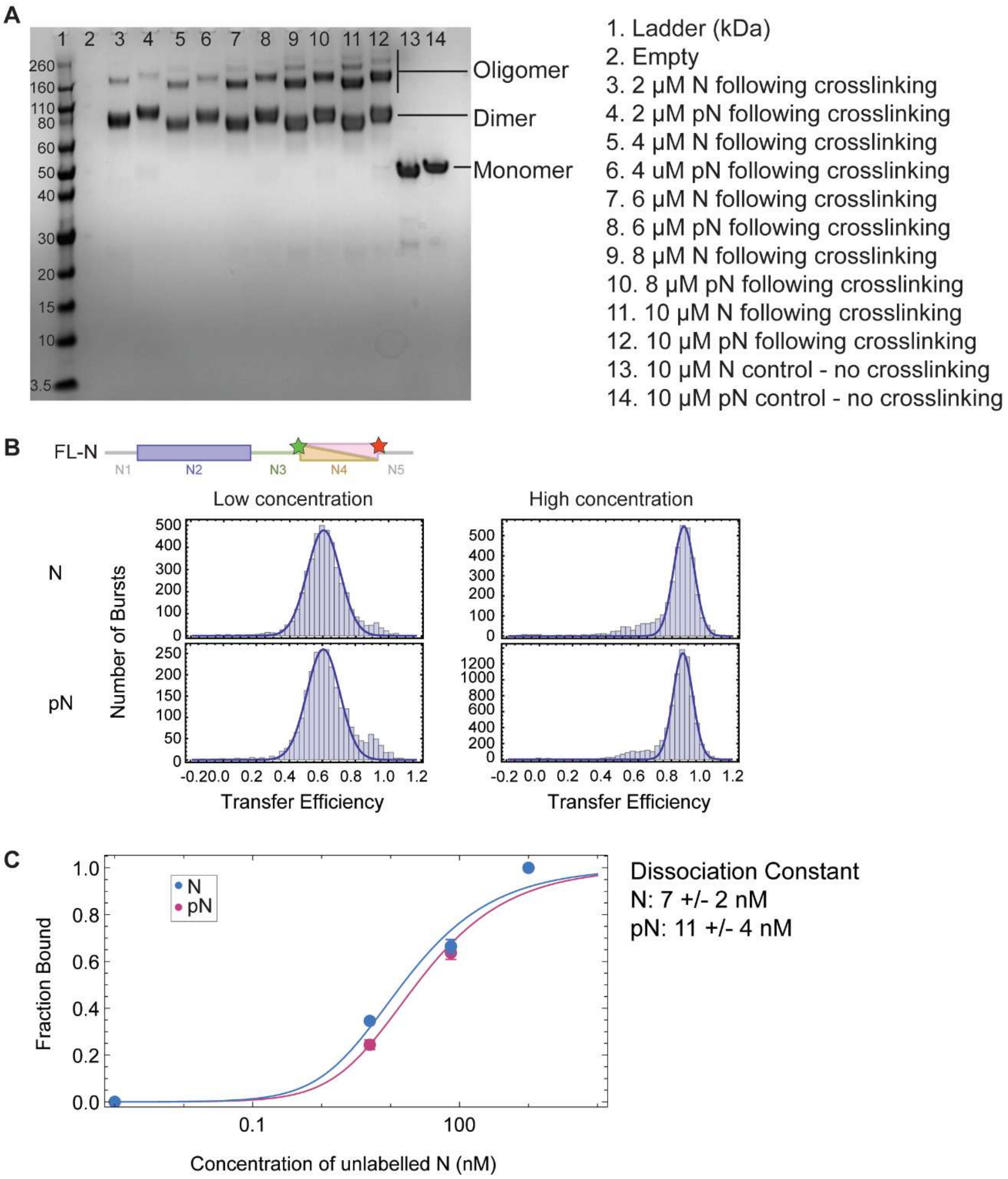


**Figure S14. Dimerization and oligomerization analysis for N vs. pN.** A) SDS-PAGE gel shows N vs. pN protein (at increasing concentrations) following chemical crosslinking. Crosslinking retains dimer / oligomer state in denaturing conditions of SDS-PAGE. N or pN protein is crosslinked using Bissulfosuccinimidyl suberate (BS3). SDS-PAGE separates oligomers by size. Increasing concentrations of protein results in a greater degree of oligomerization, but phosphorylation has little effect on oligomerization. B) Analysis of the dimerization domain conformation using smFRET. A full-length construct of N was labeled at positions 245 and 363 flanking the dimerization domain. Transfer efficiencies are similar for unmodified and phosphorylated N protein at low concentration (100 pm labeled protein, monomer regime) and high concentration (100 pm labeled + 1 μM unlabeled protein, dimer regime), indicating no shift in conformation of the dimerization domain occurs upon phosphorylation. C) Using a full-length construct that is single labeled at position 363, we measured the dimerization constant for N vs. pN protein. Binding isotherms of dimerization for N vs. pN indicate a small shift in dissociation constant (quantified to the right). Binding experiments have been analyzed accounting for the possibility of forming dimers of labeled molecules with other labeled molecules, labeled molecules with other unlabeled molecules, as well as between unlabeled molecules, according to the equation developed in Cubuk et al, 2024[2]. Error bars represent one standard deviation (±1 s.d.).

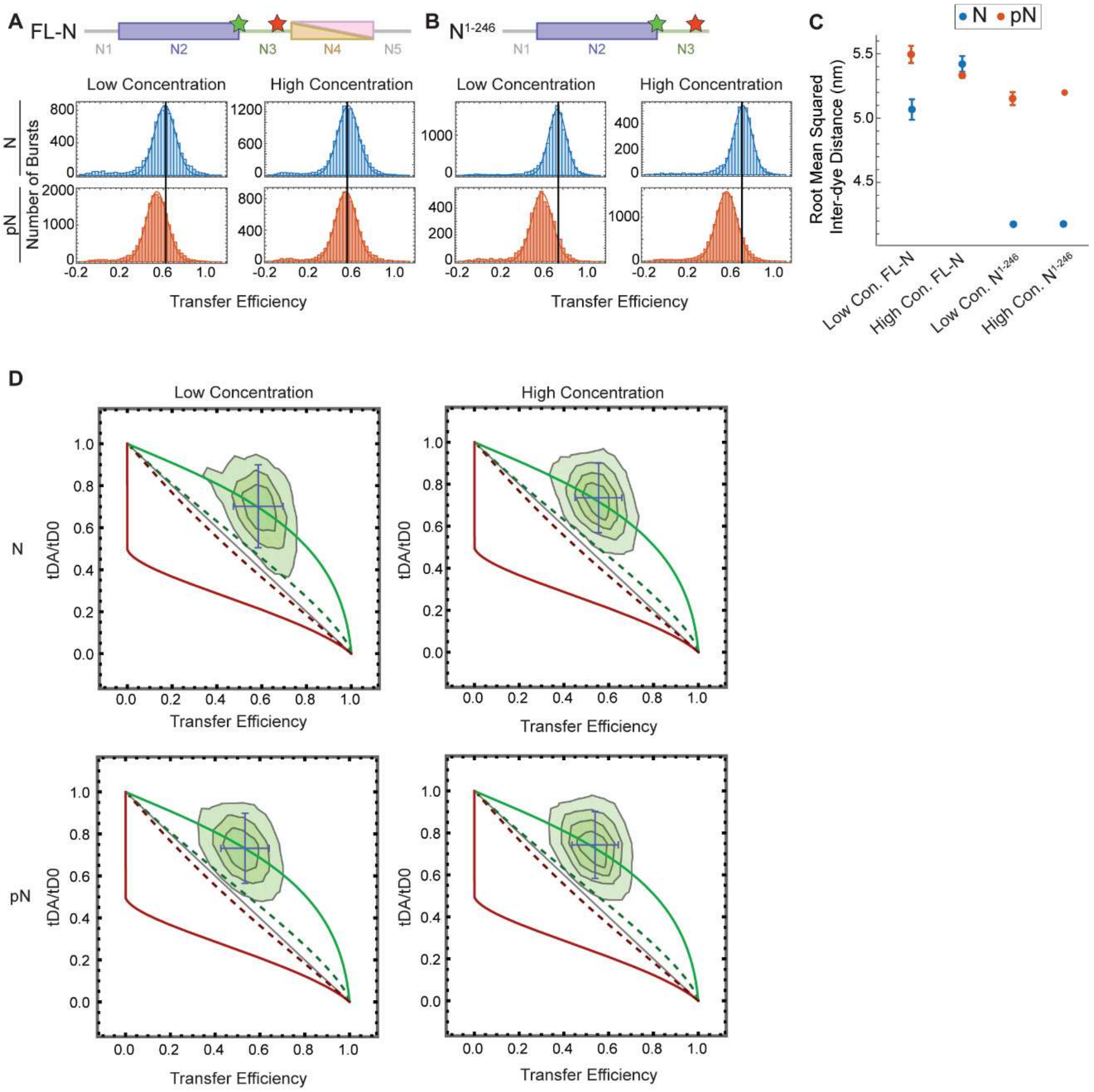


**E**

| Protein fragment | Sequence | Sequence Charge Decoration Parameter |
|---|---|---|
| N linker (N3) | GSRGGSQASSRSSSRSRNSSRNST PGSSRGTSPARMAGNGGDAALALLL LDRLNQLESKMSGKGQQQQGQTV | 2.22 |
| Phosphorylated N linker (pN3) | GSRGGSQASSRSSSRSRNSSRNST PGSSRGTSPARMAGNGGDAALALLL LDRLNQLESKMSGKGQQQQGQTV | 5.18 |

**Figure S15. Measuring the extension and dynamic nature of the linker domain.** We probed two constructs of N with fluorescent labels flanking the linker region (at residues 172 and 245): full-length N and truncated $N^{1-246}$, which lacks the dimerization and C-terminal disordered domains. We performed experiments at two protein concentrations: low concentration (100 pM labeled protein), at which N is in its monomeric form, and high concentration (100 pM labeled protein + 1 µM unlabeled protein for full-length N or 4 µM unlabeled protein for $N^{1-246}$), at which dimers form if the dimerization domain is present. We measured the distribution of transfer efficiencies for each protein construct at each concentration. A) Representative distributions of transfer efficiency for full-length N (top) and pN (bottom) at low concentration (100 pM labeled protein) and high concentration (100 pM labeled protein + 1 µM unlabeled protein) with fluorescent dyes flanking the linker region at residues 172 and 245. B) Representative distributions of transfer efficiency for $N^{1-246}$ (top) and $pN^{1-246}$ (bottom) at low concentration (100 pM labeled protein) and high concentration (100 pM labeled protein + 4 µM unlabeled protein) with fluorescent dyes flanking the linker region at residues 172 and 245. C) Root mean squared inter-dye distance obtained from the mean transfer efficiencies for unmodified and phosphorylated full-length N and $N^{1-246}$. D) Dependence of fluorescence lifetime on transfer efficiency. Comparison of fluorescence lifetimes for the full-length N in its unmodified and phosphorylated states. Grey line: linear dependence is expected for a rigid molecule. Green line: the donor lifetime (normalized by the donor lifetime in absence of acceptor: $t_{DA}/t_{D0}$) in the limit of dynamics much faster than the burst duration but slower than the fluorophore lifetime. In all cases, the populations sit near the dynamic line (green) as opposed to falling on the static line (gray), indicating the linker domain remains dynamic in both concentrations and regardless of phosphorylation status. E) Sequence charge decoration (SCD) parameter for the linker sequence prior to and following phosphorylation, which adds -2 charges to the underlined residues. A smaller SCD score indicates greater charge segregation for sequences with many positive and negative charges. SCD has been shown to be correlated with disordered proteins' radii of gyration (Rg)[3], thus indicating here that phosphorylation is expected to expand N protein's linker domain.

**A)** Mean $R_g$ for N = 5.55 nm, mean $R_g$ for N following reweighting = 5.85 nm

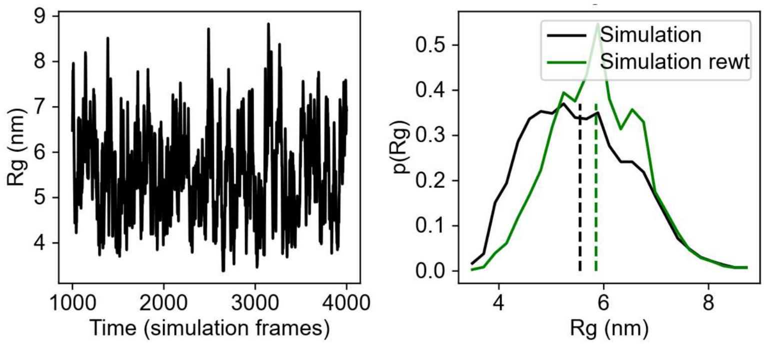


**B)** Mean $R_g$ for phosphomimetic N = 5.24 nm, mean $R_g$ for phosphomimetic N following reweighting = 5.50 nm

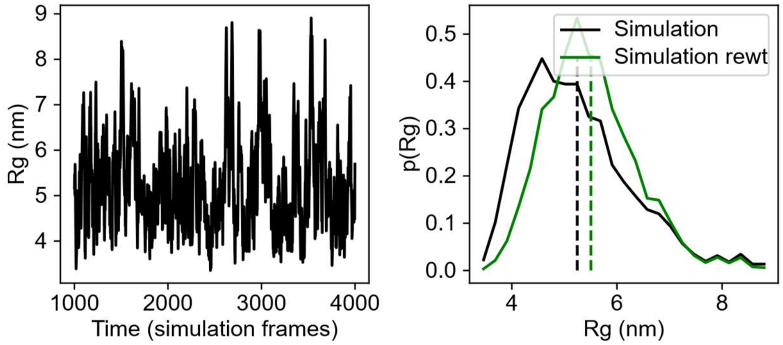


**C)**

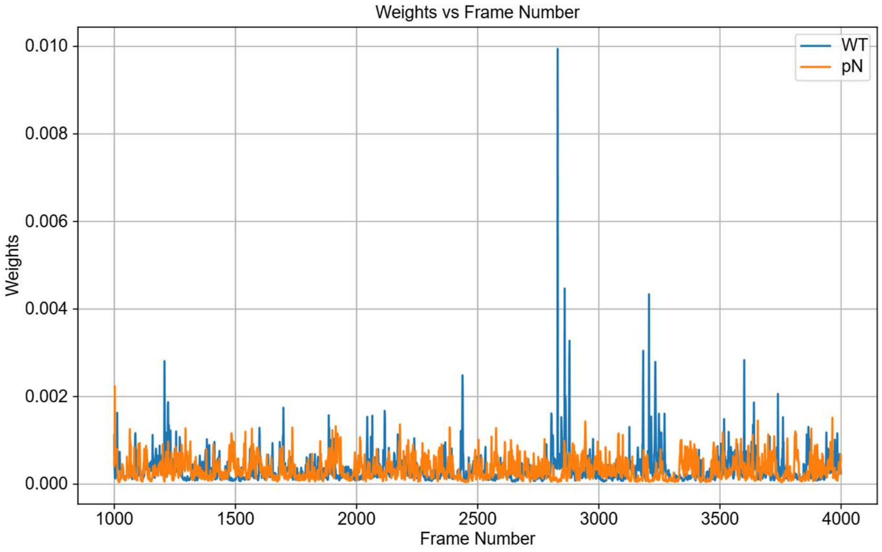


**Figure S16. Results from reweighting molecular dynamics simulation with SAXS data.** A) Left, Radius of gyration ($R_g$) from simulating N protein across frames. Right, distribution of $R_g$ from the simulation prior to (black) and after (green) reweighting.

Dashed line indicates the mean which is noted in text above. B) Left, $R_g$ from simulating phosphomimetic N protein across frames. Right, distribution of $R_g$ from the simulation prior to (black) and after (green) reweighting. Dashed line indicates the mean which is noted in text above. C) Weight of each simulation frame following reweighting for the unmodified (blue) and phosphomimetic (orange) N protein.

<table>
<tr><th>Protein and data source</th><th>Mean Radius of Gyration (nm)</th><th>Distance between residues 184-257 (nm)</th><th>Change in distance (nm)</th></tr>
<tr><td>N from original simulation</td><td>5.55</td><td>5.63</td><td rowspan="2">-0.25</td></tr>
<tr><td>Phosphomimetic N from original simulation</td><td>5.24</td><td>5.38</td></tr>
<tr><td>N after reweighting with SAXS data</td><td>5.85</td><td>5.61</td><td rowspan="2">+0.08</td></tr>
<tr><td>Phosphomimetic N after reweighting with SAXS data</td><td>5.50</td><td>5.69</td></tr>
<tr><td>N from smFRET</td><td>N/A</td><td>5.33</td><td rowspan="2">+0.16</td></tr>
<tr><td>pN from smFRET</td><td>N/A</td><td>5.49</td></tr>
</table>

**Figure S17. Comparison of distance between residues 184 and 257 spanning the N linker domain (N3).** Data was obtained from simulations prior to and following reweighting and smFRET experiments (interdye distance calculated between residues 184 and 257 based on FRET efficiencies shown in Figure S15C). Simulations approximated effect of phosphorylation by incorporating phosphomimetic mutations. Results from simulations following reweighting better capture the trend in linker expansion measured using smFRET, while still showing a decrease in radius of gyration upon phosphomimetic mutation.

### A) Intra-monomer contacts between N2 – N2 domains

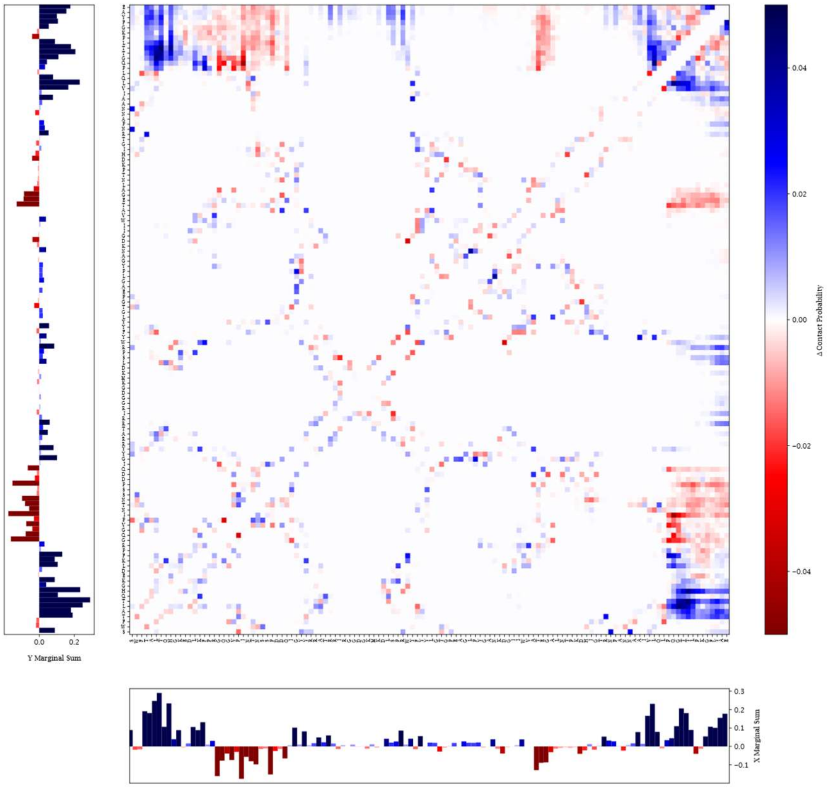


### B) Intra-monomer contacts between N3 – N3 domains

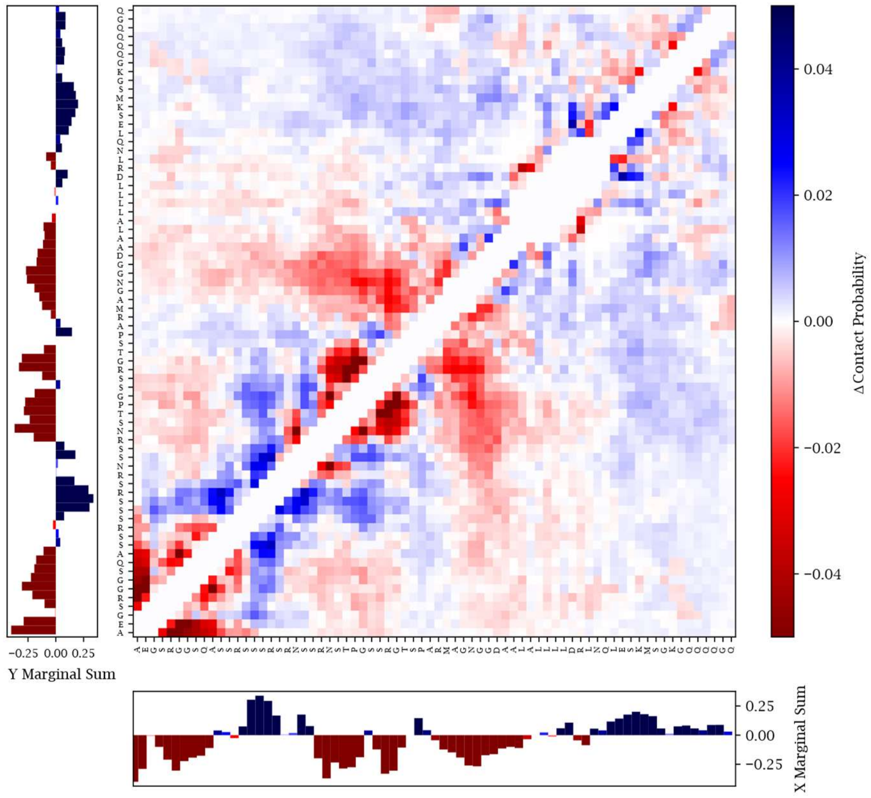

**C) Intra-monomer contacts between N3 – N4 domains**

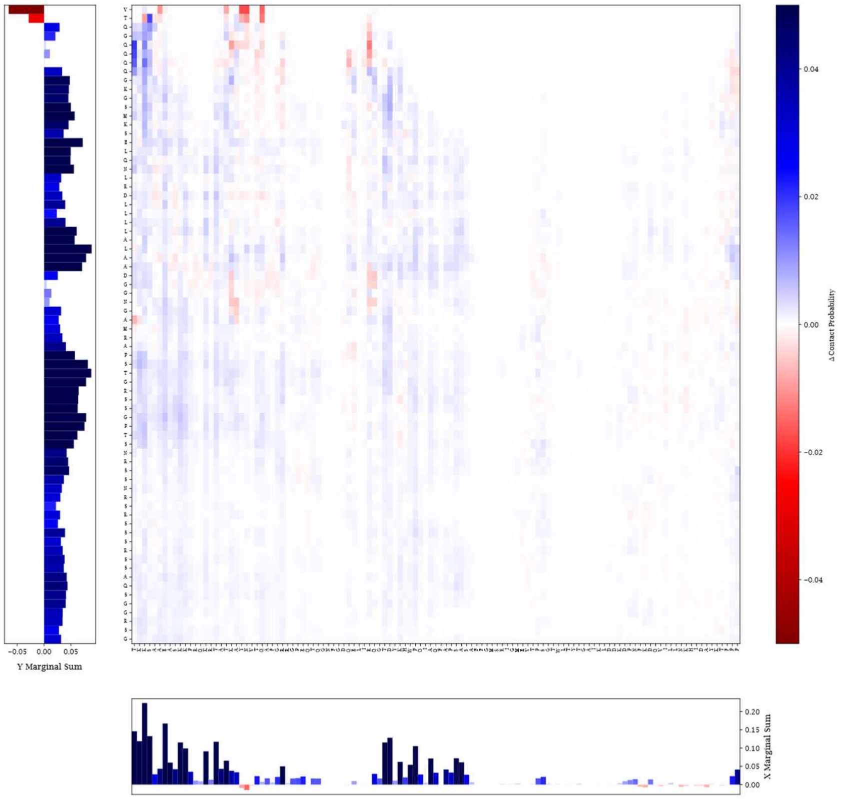

**D) Inter-monomer contacts between N3 – N2 domains**

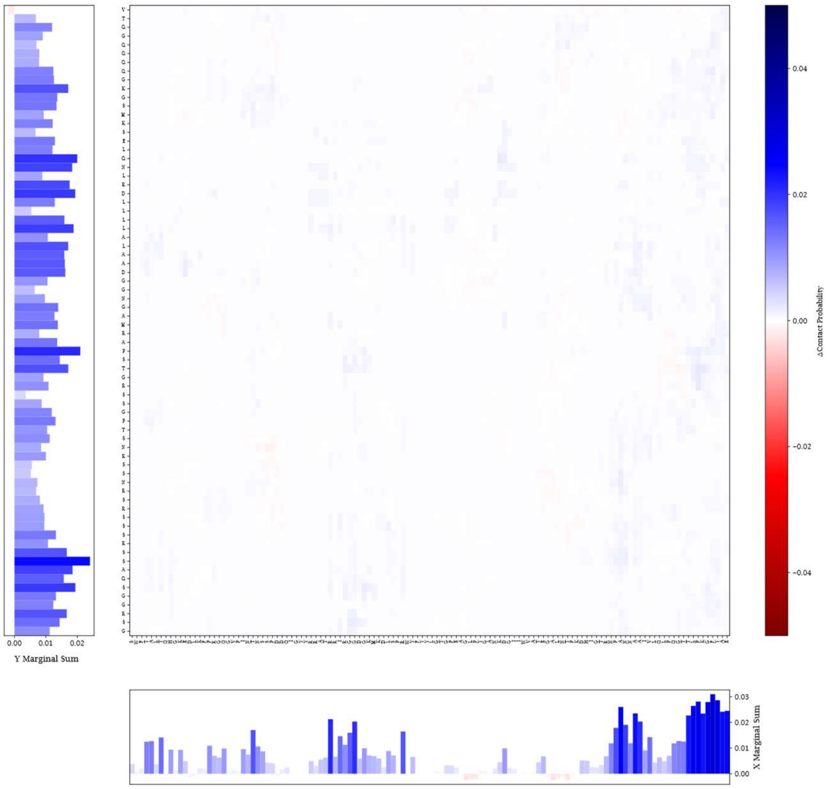

### E) Inter-monomer contacts between N3 – N3 domains

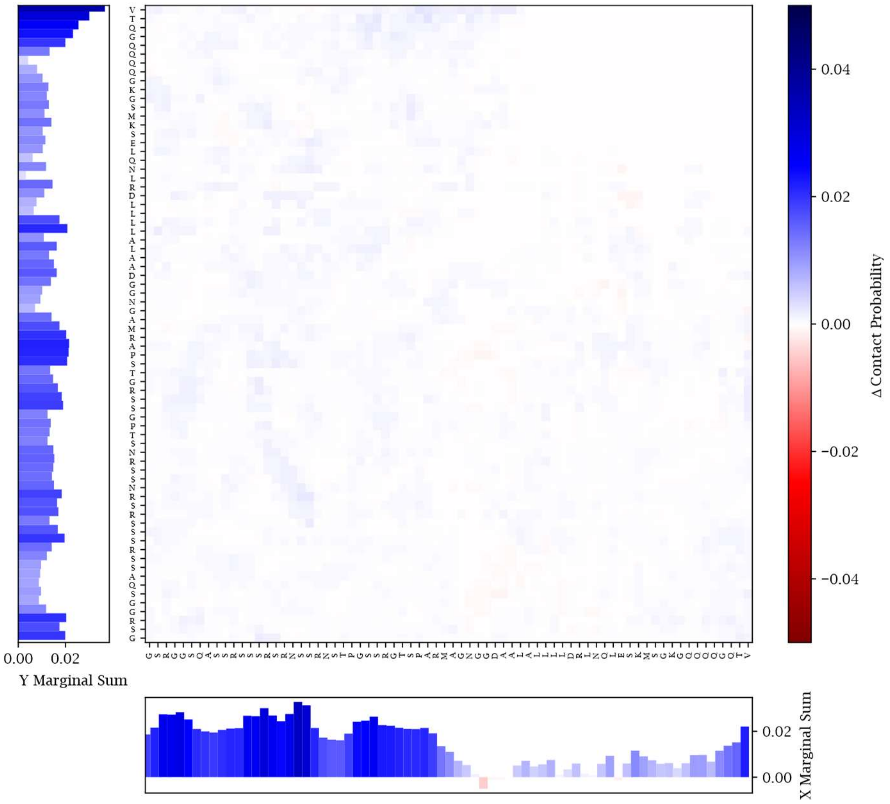


### F) Inter-monomer contacts between N3 – N4 domains

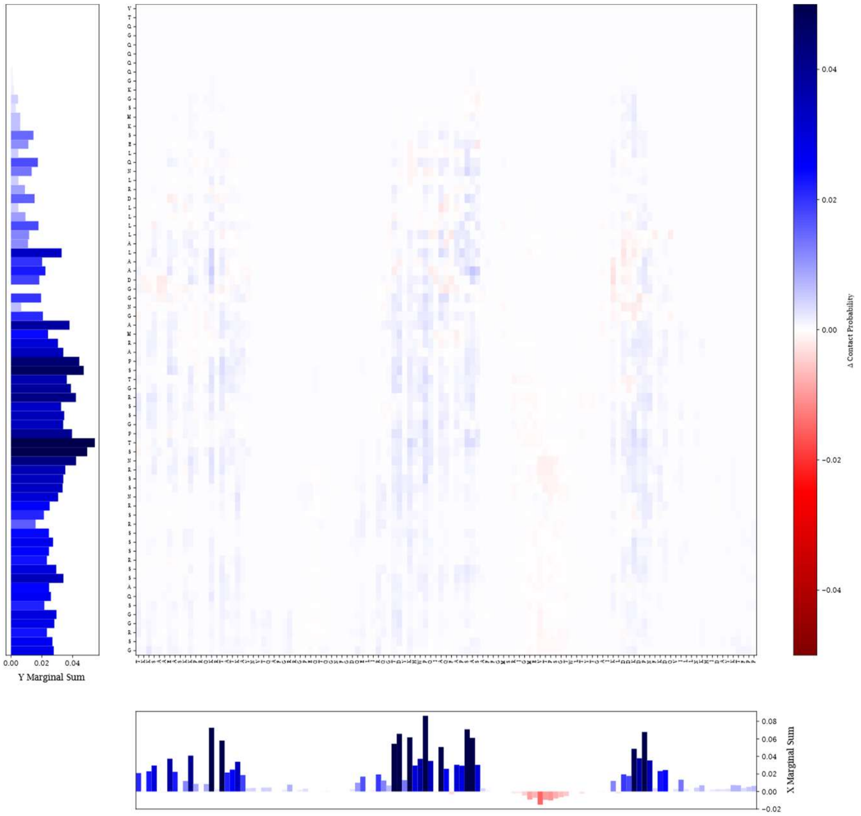

**Figure S18. Change in intra- and inter-monomer interactions between domains with phosphomimetic mutations.** A) Change in intra-monomer interactions per residue within the N2 domain following phosphomimetic mutations. B) Change in intra-monomer interactions per residue within the N3 domain following phosphomimetic mutations. C) Change in intra-monomer interactions per residue between the N3 and N4 domains following phosphomimetic mutations. D) Change in inter-monomer interactions per residue between the N2 and N3 domains following phosphomimetic mutations. E) Change in inter-monomer interactions per residue between the N3 domains following phosphomimetic mutations. F) Change in inter-monomer interactions per residue between the N3 and N4 domains following phosphomimetic mutations.

**Legends for Movies S1 to S6.**

**Movie S1 (separate file). N + 1-1000 RNA + GUVs with no membrane protein fragment.** Widefield imaging sequence of several condensates composed of N protein + 1-1000 RNA being held by optical tweezers surrounding a GUV located at the center of the field of view. One condensate is brought to the surface of the GUV and then pulled away repeatedly without causing deformation of the condensate or GUV. Video taken at 2 frames per second. Video related to Fig. 2d.

**Movie S2 (separate file). pN + 1-1000 RNA + GUVs with no membrane protein fragment.** Widefield imaging sequence of several condensates composed of pN protein + 1-1000 RNA being held by optical tweezers surrounding a GUV located at the center of the field of view. Several condensates are brought to the surface of the GUV and then pulled away without deforming the condensates or GUV. Condensates can also be held at the surface of the GUV for several seconds causing no deformation of the condensate, after which they can be pulled away from the surface. Video taken at 2 frames per second. Video related to Fig. 2d.

**Movie S3 (separate file). N + 1-1000 RNA + GUVs with Nsp3$^{1\text{-}111}$ membrane protein.** Widefield imaging sequence of two condensates composed of N protein + 1-1000 RNA being held by optical tweezers and brought to the surface of a GUV coated with Nsp3$^{1\text{-}111}$ at the center of the field of view. The condensates attach to the surface. The larger condensate falls from the focal plane after being released by the tweezer, due to gravity. Video taken at 2 frames per second. Video related to Fig. 2d.

**Movie S4 (separate file). pN + 1-1000 RNA + GUVs with Nsp3$^{1\text{-}111}$ membrane protein.** Widefield imaging sequence of several condensates composed of pN protein + 1-1000 RNA being held by optical tweezers and brought to the surface of a GUV coated with Nsp3$^{1\text{-}111}$ at the center of the field of view. The condensates wet the surface of the GUV and form a layer of protein over time. Video taken at 2 frames per second. Video related to Fig. 2d.

**Movie S5 (separate file). N + 1-1000 RNA + GUVs with M$^{104\text{-}222}$ membrane protein.** Widefield imaging sequence of several condensates composed of N protein + 1-1000 RNA being held by an optical tweezer and brought to the surface of a GUV coated with M$^{104\text{-}222}$ at the center of the field of view. The condensates attach to the surface. One condensate is wrapped by the GUV membrane over time, resulting in partial engulfment of the condensate. Video taken at 2 frames per second. Video related to Fig. 2d-e.

**Movie S6 (separate file). pN + 1-1000 RNA + GUVs with M$^{104\text{-}222}$ membrane protein.** Widefield imaging sequence of several condensates composed of pN protein + 1-1000 RNA being held by optical tweezers surrounding a GUV coated with M$^{104\text{-}222}$ located at the center of the field of view. Several condensates are brought to the surface of the GUV and then pulled away without deforming the condensates or GUV. Condensates are held at the surface of the GUV for several seconds where they can fuse with other condensates and still be pulled away from the GUV surface, causing no deformation of the condensate or GUV surface. Video taken at 2 frames per second. Video related to Fig. 2d.
